# Chromosomal and gonadal sex have differing effects on social motivation in mice

**DOI:** 10.1101/2024.10.28.620727

**Authors:** Sneha M. Chaturvedi, Simona Sarafinovska, Din Selmanovic, Katherine B. McCullough, Raylynn G. Swift, Susan E. Maloney, Joseph D. Dougherty

## Abstract

**Plain English Summary:** As our brain develops, many factors influence how we behave later in life. The brain forms differently in males and females, potentially leading to sex variation seen in many behaviors including sociability. In addition, conditions defined by differences in social behaviors, such as autism, are diagnosed more in males than females. However, researchers don’t know exactly how distinct sex factors, such as hormones and sex chromosome genes, lead to different behaviors in males and females. In this study, we used mouse models and tests of mouse behavior to explore these differences. Results show that sex hormones primarily contributed to differences in social motivation between sexes. Yet when we repeated these same assays in a mouse model of genetic liability for a human neurodevelopmental syndrome, we found that sex chromosome genes rather than sex hormones played a larger role in the behavioral consequences of impaired neurodevelopment. These insights can inform future research on the biological mechanisms of social behavior in the context of genetic liability for neurodevelopmental disorders.

**Highlights:**

- Four-core genotype mouse model crossed with MYT1L heterozygous mouse revealed independent effects of chromosomal and gonadal sex on social motivation.
- *Myt1l* haploinsufficiency was associated with increased activity in both males and females.
- While females are more active, contributions of chromosomes and gonadal hormones to this sex effect are environment dependent.
- Presence of ovaries was associated with increased measures of social seeking and orienting regardless of genotype.
- Chromosomal sex interacted with MYT1L genotype, with increased social orienting and seeking specifically in XX MYT1L heterozygous mice.

**Background:** Sex differences in brain development are thought to lead to sex variation in social behavior. Sex differences are fundamentally driven by both gonadal (i.e., hormonal) and chromosomal sex, yet little is known about the independent effects of each on social behavior. Further, mouse models of the genetic liability for the neurodevelopmental disorder MYT1L Syndrome have shown sex specific deficits in social motivation. In this study, we aimed to determine if hormonal or chromosomal sex primarily mediate the sex differences seen in mouse social behavior, both at baseline and in the context of *Myt1l* haploinsufficiency.

**Methods:** Four-core genotype (FCG) mice, which uncouple gonadal and chromosomal sex, were crossed with MYT1L heterozygous mice to create eight different groups with unique combinations of sex factors and MYT1L genotype. A total of 131 mice from all eight groups were assayed for activity and social behavior via the open field and social operant paradigms. Measures of social seeking and orienting were analyzed for main effects of chromosome, gonads, and their interactions with *Myt1l* mutation.

**Results:** The FCGxMYT1L cross revealed independent effects of both gonadal and chromosomal sex on activity and social behavior. Specifically, the presence of ovaries, and by extension the presence of ovarian hormones, increased overall activity, social seeking, and social orienting regardless of genotype. In contrast, sex chromosomes affected social behavior mainly in the MYT1L heterozygous group, with XX sex karyotype when combined with MYT1L genotype contributing to increased social orienting and seeking.

**Conclusions:** Gonadal and chromosomal sex have independent mechanisms of driving increased social motivation in females. Additionally, sex chromosomes may interact with neurodevelopmental mutations to influence sex variation in atypical social behavior.

## Background

From early development, sex differences drive distinct patterns in social motivation and social interaction in mammals[1]. In turn, social behavior plays an important role in development, as offspring learn crucial survival skills through observation and mimicry of older community members[2]. The impact of biological sex variation on the development of social behaviors is poorly understood, especially when considering the complex interaction of genetic and hormonal mechanisms contributing to sexual differentiation[3]. Sociability depends on behaviors on both sides of the interaction, including an individual’s internal drive to be social, termed social motivation, and interactions from the social partner. Social motivation is comprised of three main aspects: social seeking (orienting to a social stimulus), social reward (drive for social interaction based on benefit), and social maintenance (fostering existing social bonds)[4]. Human social interactions also include sociocultural factors, which make uncovering the biological mechanisms underlying intrinsic social motivation difficult. Therefore, rodent models have become key to understanding potentially conserved molecular and circuit contributions to typical and atypical social motivation[5].

Until recently, there were no comprehensive assays to measure social motivation in rodents. While tasks such as the three-chamber social approach assay help answer questions on overall sociability preference, they do not directly provide a quantitative measure of motivation, such as the extent to which a rodent will work for social interaction[6]. In response to this need, animal behavior experts have developed and validated a social operant protocol[7–9], designed to test two aspects of social motivation in rodents: social seeking and social orienting. Results from initial social operant cohorts in mice revealed a sex bias in social motivation, with male mice more likely to seek social interaction than female mice[9]. This supports a long history of sex variation in social behaviors in rodents.

The development of a sensitive social motivation assay has enabled the investigation of conditions in which social motivation may be altered. Autism is a neuropsychiatric condition defined by atypical social interaction[4]. The social motivation theory of autism proposes that the biological mechanisms behind autism lead to decreased social seeking, causing decreased interaction with others and therefore lower reward from social interaction. The lower social reward causes further decreased social seeking, in a behavioral loop that reinforces as children age[4]. While mice do not have all the characteristics of autism as diagnosed in humans, animal models of genetic liability are useful in investigating potential molecular mechanisms underlying autism-relevant behaviors, especially in the context of monogenic syndromes associated with autism diagnoses, as such single gene mutations that are readily recreated in mice. Such mouse studies allow for well-powered, well-controlled examination of the consequences of single gene mutations on conserved aspects of brain development and behavioral circuit functions that would be challenging to implement for patient populations.

MYT1L Syndrome is one such monogenic syndrome, with around 45% of carriers receiving an autism diagnosis[10]. This syndrome is characterized by intellectual, speech, and motor impairments in almost all carriers, and can include obesity, endocrine disruption, attention deficit hyperactivity disorder (ADHD) and epilepsy[10–13]. MYT1L syndrome can be caused by a single copy mutation in the *MYT1L* gene on chromosome 3, causing a deficiency in MYT1L protein levels[13]. MYT1L is a neuronal transcription factor essential for typical neuronal maturation[14,15]. We have previously produced a MYT1L heterozygous (Het) mouse model inspired by a local patient’s loss of function mutation[16]. In the social operant task, wildtype males showed an increase in social seeking compared to wildtype females[9,16]. Comparing the *Myt1l* mutant groups revealed a sex by genotype interaction, with male mutant mice showing a decrease in social seeking when compared to male wildtype mice. Female wildtype and mutant mice had comparable levels in social seeking. In another well-studied monogenic mouse model of autism, *Shank3b* mutation, sex specific findings were also observed. Specifically, male heterozygous and homozygous *Shank3b* mutants showed fewer social seeking behaviors than their wildtype controls, while females were unaffected[9]. Overall, this result suggests males may be more vulnerable to the effects of neurodevelopmental mutations on social behaviors.

Sex variation is complex, with sex factors such as chromosomes (XX vs. XY) and hormones (e.g., estrogen vs. testosterone) contributing to sex differences from cellular to behavioral levels. While the presence of an *SRY* gene on the Y chromosome during development determines gonadal sex, including development of testes vs. ovaries and secretion of testosterone vs. estrogen and progesterone [17], there are also effects of chromosomal sex on phenotypes independent of the action of hormones[18]. As we uncover sex differences, it is essential to untangle the effects of chromosomal and hormonal (i.e., gonadal) sex[3]. Specifically, are baseline sex differences in aspects of social motivation primarily driven directly by genes on sex chromosomes, or by hormonal signaling downstream of gonads? One way to uncouple gonadal and chromosomal sex is by using the four-core genotype (FCG) mouse model. The FCG model separates chromosomal and gonadal sex through the relocation of the *Sry* gene, responsible for testes development, to chromosome 3 instead of the Y chromosome[19]. Several labs have utilized the FCG model to untangle chromosomal and hormonal sex differences in behavior and addiction[20–23]. The results from these studies have found strong evidence of the role of gonadal hormones in sexually dimorphic behavior, replicating and further validating past gonadectomy studies[24]. More interestingly, many studies found an independent role for X chromosome genes on sex variation in behavior, commonly through X-gene number (or dosage) or parental imprint[21]. Detangling the relative effects of sex hormones and sex chromosomes allows researchers to focus subsequent resources on mechanisms that drive behavioral sex variation[20].

No currently published studies have combined the FCG model with a model of altered neurodevelopment. Therefore, in addition to understanding typical sex variation in social behavior, we aim to understand whether the sex by genotype interaction in our MYT1L haploinsufficiency model[16] is due to either chromosomal or gonadal sex. A high percentage of patients with MYT1L Syndrome have endocrine issues, suggesting potential dysregulation in the hypothalamic-pituitary axis which can lead to dysregulated sex hormone variation. On the other hand, MYT1L is a transcription factor and could interact with genes on the X or Y chromosomes. Therefore, to focus future experiments on either a hormonal or transcriptional mechanism for *Myt1l* mutation’s effects on behavior, we first aimed to establish if the sex by genotype interaction seen in *Myt1l* mutants was driven by chromosomal or gonadal sex. Thus, we crossed the FCG model with our MYT1L model, producing eight different groups, with four different sex combinations (XXF [ovaries], XXM [testes], XYF [ovaries], XYM [testes], split by MYT1L wildtype [WT] and mutant genotypes). All groups were tested in the open field and social operant assays to investigate sex and genotype effects on activity and social motivation. Results from both behavior assays demonstrated increased activity driven by presence of ovaries. Similarly, the presence of ovaries increased social seeking and orienting in the social operant assay. Interestingly, XX sex karyotype also increased social behaviors, but only in the context of *Myt1l* haploinsufficiency. These results highlight the potential role of X chromosomal genes in mediating sex variation, especially during altered neurodevelopment.

## Methods

### Animal Models

All procedures using mice were approved by the Institutional Care and Use Committee at Washington University School of Medicine. Mice were bred and maintained in the vivarium at McDonnell Medical Sciences Building at Washington University in St. Louis in static (28.5 x 17.5 x 12 cm) translucent plastic cages with corncob bedding and *ad libitum* access to standard lab diet and water. Animals were exposed to 12/12-hour light/dark cycle, at 20-22°C and 50% relative humidity. Breeding pairs for experimental cohorts were comprised of female *Myt1l* Het (JAX Stock No. 036428) and male XY*^Sry-^* mice on a C57BL/6J background (JAX Stock No.010905) to generate eight experimental groups (**Fig 1A**). Sample sizes are listed in Figure 1B. Animals were weaned at P21, and group-housed by gonadal sex and MYT1L genotype. C57BL/6J mice (JAX Stock No 000664) were used as social partners during behavioral testing.

**Figure 1.**
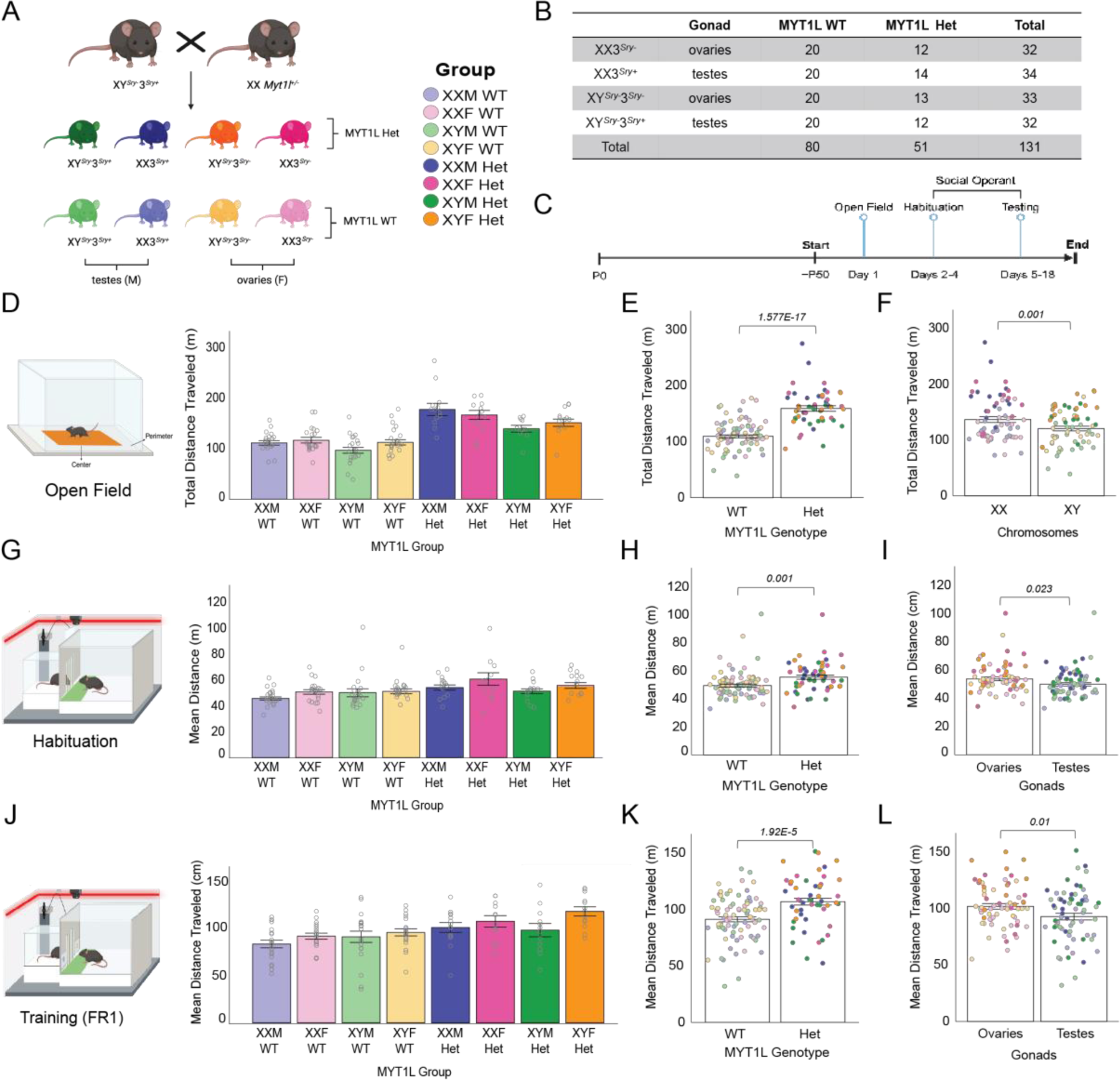
Myt1l heterozygous mice are more active than Myt1l wildtype in both social operant and open field tasks. **A)** Four core mouse model crossed with *Myt1l^+/-^* female to create eight separate groups with unique combinations of *Myt1l* variant, sex chromosome karyotype, and gonadal sex**. B)** Breakdown of sample number, with a range of 12-20 mice per group. **C)** Experimental timeline demonstrating start of behavioral assays at ∼P50 [P45-P61] with open field, followed by habituation and social operant. **D)** Total distance traveled by each group in the open field (OF) task. Darker colors designate MYT1L Het, lighter colors designate Myt1l WT. **E)** Myt1l Het mice traveled a significantly greater distance than wildtype in OF. **F)** Mice with XX chromosomes traveled a significantly greater distance than mice with XY chromosomes in OF. **G)** Mean distance traveled per day by each group in the social operant (Soc Op) task during the habituation period. During habituation, door is maintained open, and nose poke panels are removed. **H)** Myt1l Het mice traveled a greater distance per day than MYT1L WT in Soc Op habituation. **I)** Mice with ovaries traveled a greater distance than mice with testes in Soc Op habituation. **J)** Mean distance traveled per day by each group in Soc Op during the training period fixed ratio 1 (FR1). **K)** Myt1l Het mice traveled a significantly greater distance than Myt1l WT in Soc Op FR1. **L)** Mice with ovaries traveled a significantly greater distance than mice with testes in Soc Op FR1. Error bars in all panels indicate standard error of the mean (SEM).

### Genotyping

*Myt1l* genotyping of breeders and experimental litters before behavioral assays were conducted with allele specific PCR using *Myt1l* mutant and control primers [16]. Four-core genotype (FCG) was determined with allele specific PCR using an established protocol by The Jackson Laboratory (Protocol 5990: Standard PCR Assay – Tg(Sry)Eicher). Genotypes of all experimental mice were reconfirmed at the end of the experiment.

### Behavior Testing

For behavioral analysis, five batches of 131 mice total (**Fig 1B**) were used to assess activity and social motivation. All tasks were run by a female experimenter, during the light phase. Mice were handled for 3 days prior to starting the first behavioral task and the tails of mice in were marked with a non-toxic, permanent marker regularly to easily distinguish mice during testing. Male gonadal and female gonadal cages were separated in the testing room to avoid olfactory cue influence on behavior. Testing orders were randomly counterbalanced for group across apparatuses and trials. Testing began around P50 (P45 – P61) for all animals with open field followed by the social operant assay (**Fig 1C**).

### Open field

Locomotor activity was measured to assess activity, exploration, and anxiety-like levels in the open field assay similar to our previous work[25]. Briefly, each mouse was recorded individually for a 1-hr period in a white matte acrylic apparatus measuring 40×40 cm, inside a custom sound-attenuating chamber (70.5 × 50.5 × 60 cm), with red 9 lux illumination (LED Color-Changing Flex Ribbon Lights, Commercial Electric). A CCTV camera (SuperCircuits) controlled by ANY-maze software (Stoelting Co.; http://www.anymaze.co.uk/index.htm) tracked each mouse within the apparatus to quantify distance traveled, time in, and entries into pre-established center/perimeter zones. The apparatus was cleaned between animals with a 0.02% chlorhexidine diacetate solution (Nolvasan, Zoetis).

### Social operant

Social motivation, specifically measures of social reward seeking and social orienting as defined below, was evaluated beginning one day after open field using a social operant task adapted from previous methods[9]). Standard operant chambers (Med Associates) enclosed in sound-attenuating chambers (Med Associates) were modified to contain a clear acrylic box used for a ‘stimulus chamber’ (10.2 × 10.2 × 18.4 cm; Amac box, The Container Store) attached to the operant chamber, separated by a raisable door (10.2 × 6 cm) and stainless-steel bars (6mm spacing), flanked by nosepoke holes (**Fig. 2A**). The door was connected to an Arduino (UNO R3 Board ATmega328P) controlled by Med Associates software. Operant chambers and stimulus chambers were designated for males or females throughout the experiment, defined by gonadal sex. The operant chambers were cleaned with 70% ethanol and the stimulus chambers were cleaned with 0.02% chlorhexidine diacetate solution (Nolvasan, Zoetis) between animals.

**Figure 2.**
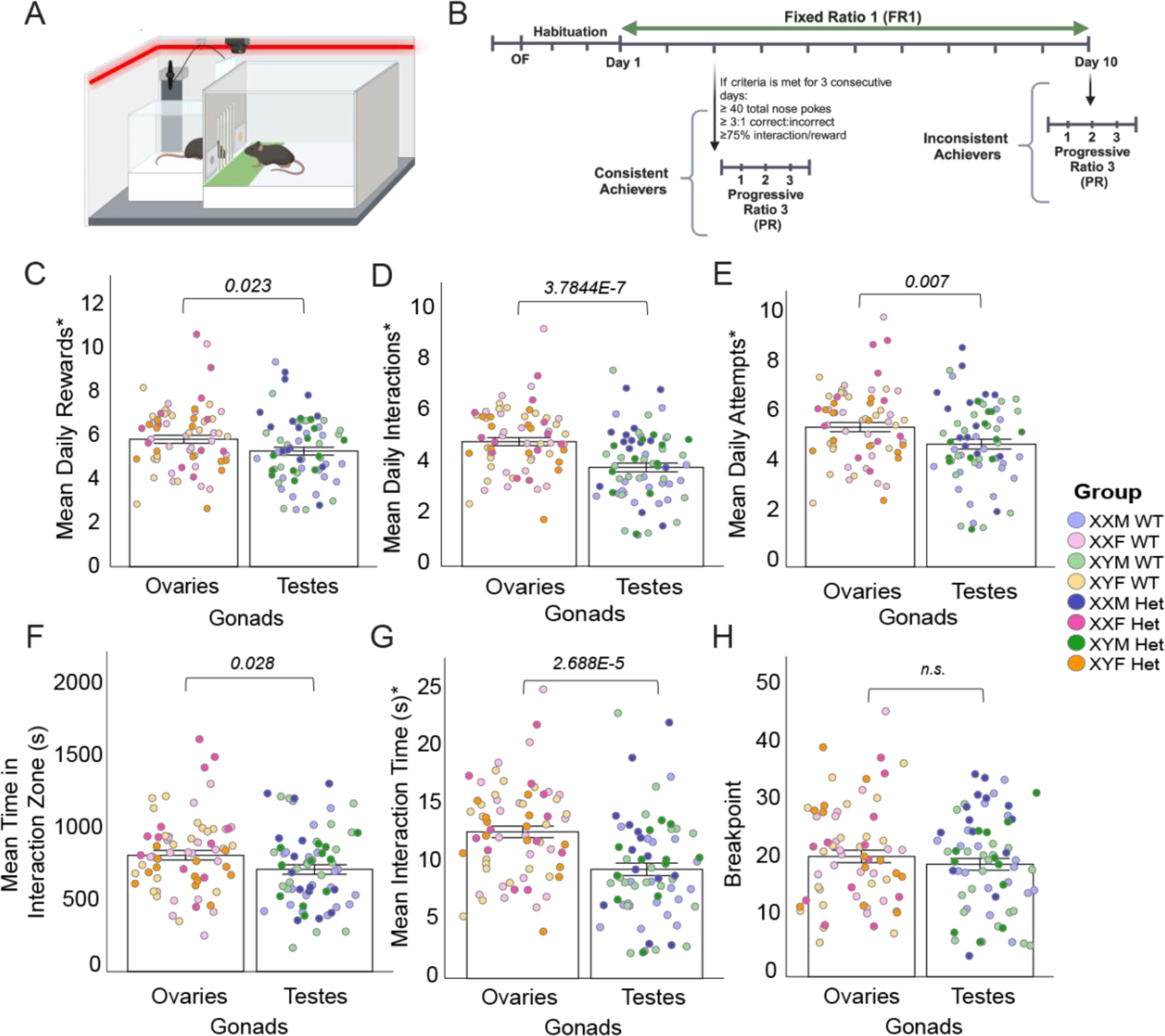
Mice with ovaries perform higher in variables of social seeking and orienting than mice with testes. **A)** Diagram of social operant chamber, with active nosepoke in yellow and door open to allow interaction between stimulus (left) and test (r ight) mouse. Right line designates LED light strip. Green rectangle indicates “Interaction Zone”. **B)** Timeline of behavioral assays, including two days of open field, two days of habituation, and 3-10 days of fixed ratio 1 (FR1). Mice drop from FR1 to Progressive Ration (PR) once they meet three consecutive days of conditioning criteria. PR continues for three total days. **C)** Mice with ovaries receive a greater number of rewards (open door) per day compared to mice with testes. **D)** Mice with ovaries have a greater number of interactions (both stim and test mouse) per day compared to mice with testes. **E)** Mice with ovaries attempt to interact significantly more than mice with testes, per day. **F)** Mice with ovaries spend more time in the interaction zone, regardless of reward status or location of the stimulus mouse. **G)** Mice with ovaries tend to have longer interaction times with both the stimulus and test mouse in the interaction zone, compared to mice with testes. **H)** Breakpoint, or the highest number of continuous nosepokes a test mouse will complete for one reward, showed no different in regard to ovaries or chromosomes (*not shown)*. For all panels, error bars indicate SEM. Asterisk (*) indicates variables that underwent square root transformation to normalize data distribution.

The operant paradigm comprised habituation (Hab), training (fixed ratio 1; FR1), and testing (progressive ratio; PR) trials (**Fig. 2B**). For all trials, gonadal-sex- and age-matched, novel C57BL/6J mice served as stimulus mice. The stimulus mice were loaded into and removed from the stimulus chambers prior to the placement and after removal of the experimental mice into the operant chambers, respectively, to prevent the experimental animals from being in the chambers without a stimulus partner. Habituation consisted of a 30-minute trial on each of two consecutive days, during which the door remained opened, and the nosepoke holes were blocked by panels to prevent any nose-poking prior to training. Subsequent training days consisted of 1-hr trials during which the fixed ratio 1 reinforcement schedule was used to reward the mouse with a 12 s social interaction opportunity following one correct nosepoke. During the 12 s reward period, any additional correct nosepokes did not result in another reward. Task achievement criteria were displaying a) at least 40 correct nosepokes, b) a 3:1 correct: incorrect nosepoke hole ratio, and c) at least 75% of rewards including a social interaction (defined as both experimental and stimulus mice in their respective social interaction zones simultaneously for at least 1 s of the reward). After three consecutive days of showing achievement of achievement criteria resulted in the mouse attaining “consistent achiever” status and moving on to the final testing portion. Ten days of FR1 without reaching three consecutive days of criteria resulted in “inconsistent achiever” status. For the final testing portion, to measure the breakpoint, or maximum nosepokes or effort the animal would exhibit for a social reward, the progressive ratio 3 (PR3) reinforcement schedule was used to reward the mouse with a 12 second social interaction opportunity following a progressive increase in required correct nosepokes by 3 (e.g., 3, 6, 9, 12, etc.), which lasted for 3 consecutive days. Assay differences from our previously published protocol included chamber illumination with a red strip light (LED Color-Changing Flex Ribbon Lights, Commercial Electric) to achieve 75-80 lux of red light, removal of a fixed ratio 3 interval, and extending the progressive ratio testing to 3 days. A detailed description of Hab, FR1, and PR3 experimental outcomes can be found in **Supplementary Table 1.**

### Statistical Analysis

Statistical analyses and graph plotting were performed using IBM SPSS Statistics (v.26) and GraphPad Prism (v.8.2.1). Biorender.com was used for components of Figures 1, 2, and S1. All variables collected from habituation (Hab) and FR1 were averaged over number of days, to account for varying number of days in the assay per mouse. Prior to analyses, data was screened for missing values and fit of distributions with assumptions underlying univariate analysis. This included the Shapiro-Wilk test on *z*-score-transformed data and qqplot investigations for normality, Levene’s test for homogeneity of variance, and boxplot and *z*-score (±3.29) investigation for identification of influential outliers. One animal was excluded for analysis of Total Distance (OF) and Perimeter Distance (OF) (z-score > 3.21). Means and standard errors were computed for each measure. For data that did not fit normal univariate assumptions, transformations were applied. The following measures required square root transformation to normalize data distribution: Time in Center (OF), Time in Perimeter (OF), Mean Time per Center Visit (OF), Total Time in Interaction Zone (Soc Op Hab), Daily Rewards (Soc Op FR1), Daily Correct Nosepokes (Soc Op FR1), Daily Attempts (Soc Op FR1), Daily Interactions (Soc Op FR1), and Interaction Time (Soc Op FR1). Entries into Interaction Zone (Soc Op Hab) required natural logarithmic transformation to normalize data distribution. Analysis of variance (ANOVA) was used to analyze data where appropriate. First model iteration included the following fixed effects:

Chromosomes, Gonads, MYT1L, Chromosomes*Gonads, Chromosomes*MYT1L, Gonads*MYT1L, Chromosomes*Gonads*MYT1L.

Subsequent ANOVA iterations removed any non-significant interactions, until most parsimonious model was developed. Simple main effects tests (e.g., T-tests) were used to dissect significant interactions *post hoc*. Multiple pairwise comparisons were subjected to Bonferroni correction or Dunnett correction. In three cases (Total Distance, Perimeter Distance, and Mean Time per Center visit), transformation and outlier removal were not sufficient to completely normalize data, although distributions were very close. For these variables, non-parametric testing was used to confirm results from ANOVA testing. Batch effect was analyzed separately through ANOVA for an overall main effect. If there was a significant main effect, Batch, Batch*MYT1L, Batch*Gonads, and Batch*Chromosomes were added to the simplified univariate model. For batch data that did not fit normal univariate assumptions, non-parametric tests were used to determine main effects. Chi-square or Fisher’s exact tests were used to assess *Myt1l* mutation and gonadal/ chromosomal sex association with consistent vs inconsistent achiever status. The critical alpha value for all analyses was p < 0.05. The datasets generated and analyzed during the current study are available from the corresponding author upon reasonable request. All statistical data can be found in **Supplementary Table 2**.

### Mega-Analysis

Our mega-analysis included social operant experiments conducted locally and published between 2019 and 2023[9,16,26] (**Supplementary Table 3)**. In total, 143 control mice were included from seven cohorts. Daily Rewards, Total Time in the Interaction Zone, and Distance from all cohorts underwent statistical testing for effects of sex, as described above. Two animals were excluded for analysis of Daily Rewards (z-score > 3.6 and z-score < −3.29) and one animal was excluded for analysis of Total Time in the Interaction Zone (z-score > 3.7). Daily Rewards and Total Time in the Interaction Zone were square root transformed before analysis. Analysis of variance (ANOVA) was used to analyze for sex, age, and cohort interactions. Mice were grouped into three age buckets for ANOVA testing: P44-62, P66-86, and P90-110. First model iteration included the following fixed effects.

Cohort Age Sex Cohort*Age Age*Sex Sex*Cohort Cohort*Sex*Age, with cohort as a random variable. Subsequent ANOVA iterations removed any non-significant interactions and main effects, until the most parsimonious model was developed. Fixed variables were Sex and Age. Cohort was entered as a random variable. This analysis was conducted for control mice of all cohorts together. All statistical details can be found in **Supplementary Table 4**

## Results

### *Myt1l* mutants are more hyperactive regardless of sex factor combination

To isolate sex chromosome and gonadal influences on the sex bias in social motivation previously seen in our MYT1L heterozygous (Het) mice[16], we crossed a FCG XY*^Sry-^* male with a *Myt1* Het female to create eight different groups (**Fig.1A)**. Inheritance of the mutated *Myt1l* allele was less than the 50% expected, contrary to prior observations, suggesting subpar viability or fertility of mutant eggs or embryos in this cross (**Fig.S1**). Thus, final group sizes ranged from 12 to 20 (**Fig.1B)**. The eight combinations allow us to separate effects due to gonads, chromosomes, and their interaction with genotype. Once mice reached young adulthood (∼P50), we first tested them with the open field assay to assess activity levels. The next day, all groups began the social operant protocol (**Fig.1C**) to examine social phenotypes.

In the open field, MYT1L Het mice traveled a significantly greater distance overall when compared to MYT1L WT mice (**Fig.1D-E**), replicating previous findings[16]. When comparing chromosomal sex, XX mice traveled a larger distance than XY mice (**Fig. 1F**), regardless of MYT1L genotype or gonadal sex. There were no main effects of gonadal sex on total distance traveled in this assay. Although our primary goal of the open field was to test locomotion and activity, it can also be used to look at anxiety-like behavior by comparing how much time a mouse spends in the center (higher risk) vs. the perimeter (lower risk). Breaking down total distance between the predefined “center” and “perimeter” shows the same effects, with increased activity in both center and perimeter in MYT1L Het and in XX mice, independently **(Fig.S1).** Entries into the center and into the perimeter were dependent only on genotype, with MYT1L Het mice entering both areas significantly more than MYT1L WT mice **(Fig.S1)**, a result that aligns with the increased activity seen in total distance. Chromosomes were the only factor to influence time spent in each area, with XY mice more likely to spend time in the center and therefore potentially less avoidant than XX mice (**Fig.S1)**. In addition, XY mice and MYT1L Het mice had longer visits in the center on average when compared to XX and MYT1L WT mice.

Although the social operant assay primarily tests aspects of social motivation, locomotor activity as measured by distance traveled is also tracked. Therefore, we also compared distance traveled results between the open field and social operant assays. Similar to open field, MYT1L Het mice in social operant during both the habituation and testing period traveled a significantly greater distance overall when compared to MYT1L WT mice **(Fig.1G-H**, **Fig.1J-K)**. However, chromosomal sex showed no effect on distance traveled in this assay. During habituation and testing ovaries were also associated with greater distance traveled **(Fig.1I**, **Fig.1L)**, independent of MYT1L genotype.

### Ovaries are associated with increased social motivation independent of MYT1L expression

After open field testing, groups were habituated for two days to the social operant chamber with open-door access to a novel stimulus animal in the stimulus chamber (**Fig.1C**, **Fig.2A-B**). Over the course of testing, experimental mice were matched with stimulus mice of the same age and gonadal sex, with a novel stimulus mouse each day. All groups learned the operant tasks successfully, as seen from correct nosepokes being consistently higher than incorrect nosepokes across the entire testing period **(Fig.S2)**. Mice who reached criteria (described in *Methods*) are designated as “consistent achievers” while those that did not achieve criteria are designated “inconsistent achievers”. There were no significant effects from chromosomes, gonads, or MYT1L genotype on the number of consistent achievers in each group or an effect on the day criteria was met amongst consistent achievers (**Fig.S2**).

Measures defined as assessing social reward-seeking include number of rewards a mouse solicited during FR1 and PR3 stages, and PR3 breakpoint (i.e., the maximum number of pokes a mouse will exhibit for a single reward). Social orienting is interpreted through analyzing total time spent interacting with the stimulus mouse when available, alongside other variables (**Table S1)**. When analyzing the mean daily rewards between groups, gonadal sex had a significant main effect, with ovaries associated with a higher number of rewards on average per day (**Fig.2C**). This effect was driven primarily by the WT mice **(Fig.S3)**. In contrast, neither chromosomal sex nor MYT1L genotype showed any main effect on daily rewards (**Table S2**). Correct nosepokes, which unlike rewards can be obtained during the reward interval, also exhibited a gonadal effect, with ovaries associated with a higher number of correct nosepokes per day (**Table S2**). This reinforces the role of ovaries, and by extension hormones such as estrogen, having the chief effect on increasing social motivation in females.

The overall gonadal effect in outcomes associated with social seeking was also seen in measures of social orienting. The daily number of interactions (both test and stimulus mouse at the door) and attempts (only test mouse at the door) demonstrated a strong gonadal influence, with ovaries associated with a greater daily mean of interactions and attempts (**Fig.2D-E).** Along with the number of interactions, time spent in the interaction zone (test mouse only) and interaction time (test and stimulus mouse) showed a strong gonadal effect, with ovaries associated with a longer time spent in the interaction zone (**Fig.2F-G).** It is of note that when we analyze only the consistent achievers, this gonadal effect is still seen in mean daily interactions and mean daily total interaction time (**Table S3**). There were no main effects of chromosomes or gonads when examining PR breakpoint, implying males and females have a similar upper limit of social motivation (**Fig.2H**). Overall, analyzing specifically for gonadal effects, our results indicate females in this cohort have a higher social motivation due to sex hormones, and this sex effect persists regardless of *Myt1l* mutation. Analysis of the entire cohort supported an overall effect driving higher sociability in females, deviating from previous data showing higher sociability in males[9,16], which motivated a mega-analysis of social operant behavior described at the end of the results section.

### Chromosomal sex interacts with *Myt1l* mutation to increase sociability in XX mice

Unlike gonadal sex, chromosomal sex did not show an overall main effect on metrics of social seeking or social orienting. However, when accounting for MYT1L genotype, there was a significant chromosomal sex by genotype interaction, where two X chromosomes drove higher number of rewards and correct nosepokes only when comparing XX and XY MYT1L Het mice (**Fig.3A-B)**. Additionally, number of rewards and correct nosepokes showed an effect of MYT1L genotype within the XX chromosome group, with Het showing increased social seeking compared to WT. Therefore, sex chromosomes interacted with *Myt1l* genotype to affect social seeking, both within the MYT1L Het group and the XX karyotype group.

**Figure 3:**
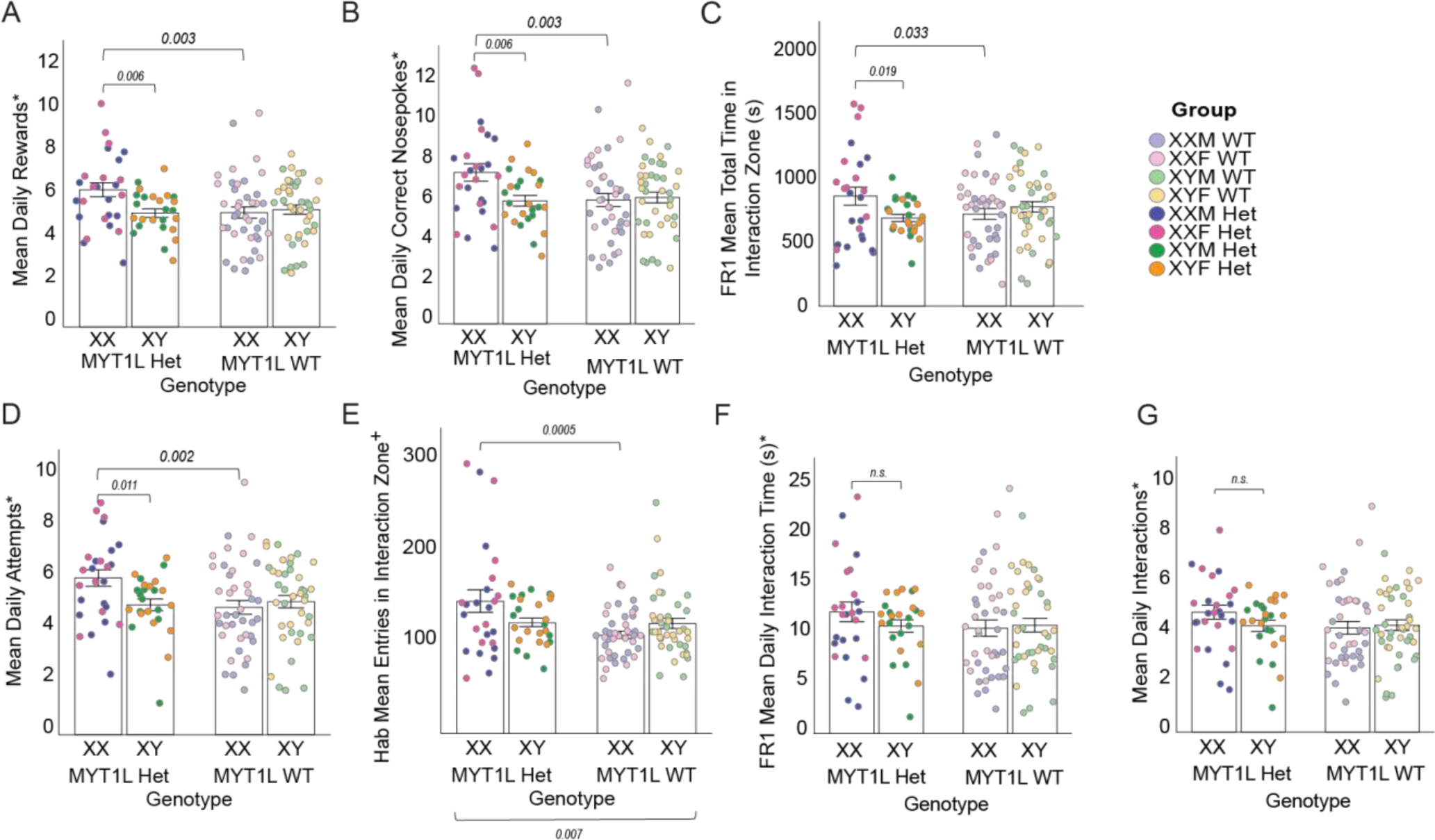
XX karyotype significantly interacts with MYT1L genotype to cause increased social motivation in females. **A)** MYT1L Het mice with XX karyotype have significantly greater mean daily rewards compared to all other groups. **B)** MYT1L Het mice with XX karyotype have significantly more correct nosepokes per day compared to all other groups. **C)** MYT1L Het mice with XX karyotype spend more time on average in the interaction zone than all other groups during FR1. **D)** MYT1L Het mice with XX karyotype make significantly more daily attempts to interact than all other groups. **E)** During habituation, MYT1L Het mice enter the interaction zone significantly more than MYT1L Het mice. Chromosomal sex and MY1L genotype interact, as XX MYT1L Het mice enter the interaction zone significantly more than XX MYT1L WT mice. **F)** MYT1L genotype and chromosomal sex show no effect on daily interaction time during FR1 (require both stimulus and test mice). **G)** MYT1L genotype and chromosomal sex show no effect on number of daily interactions (require both stimulus and test mice). For all panels, error bars indicate SEM. Asterisk (*) indicates variables that underwent square root transformation to normalize data distribution.

When looking at variables of social orienting that only consider the test mouse (total time in the interaction zone and attempts) during FR1, we continue to see the chromosomal sex by genotype interaction **(Fig.3C-D).** XX MYT1L Het mice spent more time in the interaction zone compared to XY MYT1L Het mice. Additionally, XX MYT1L Het mice spent more time in the interaction zone than XX MYT1L WT mice. A similar pattern was seen when analyzing attempts, with chromosomal sex by genotype interactions within both the MYT1L Het and XX karyotype groups. When time in the interaction zone is measured during the 2-day Hab period, this interaction is not significant with primary variation due to batch effects **(Fig.S3)**. Entries into interaction zone by the test mouse during both the Hab and FR1 period were dependent on MYT1L genotype, with MYT1L Het mice entering the interaction zone significantly more than MYT1L WT mice **(Fig.S3)**. Entries into interaction zone during the habituation period showed a chromosome by MYT1L genotype interaction within the MYT1L Het group, with XX MYT1L Het spending more time in the interaction zone than XY MYT1L Het **(Fig.3E)**. This was not seen in MYT1L WT groups, and no interaction was seen in the entries to the interaction zone during FR1 **(Fig.S3)**. Entries into the interaction zone also showed a significant interaction effect of MYT1L genotype and the XX karyotype group, with XX MYT1L Het entering the interaction zone significantly more times than the XX MYT1L WT group.

Focusing analysis on consistent achievers during FR1, this chromosomal sex by genotype interaction is maintained in rewards, total time in the interaction zone, and attempts **(Fig.S3)**. MYT1L Het mice with XX chromosomes performed higher than other groups, independent of gonadal sex. In measures that depend on both the stimulus and test mouse (total interaction time and daily interactions), the chromosomal sex by genotype interaction is no longer significant **(Fig.3F-G),** potentially due to variation in stimulus mouse behavior.

### Mega-analysis of seven social operant cohorts reveals complex sex bias in sociability

Since past published social operant results from our lab[9,16] have shown greater social motivation in males compared to females, but the data from our MYT1L WT here showed strong evidence of higher social motivation in females, we sought to better understand which outcome was the most consistent across studies. To evaluate this, we conducted a combined analysis of seven separate social operant experiments conducted between 2019 and 2023 to determine how reliable the baseline sex difference in wildtype animals replicated amongst cohorts. We also explore age and other factors that might explain differences across cohorts. **Supplementary Table 4** includes a breakdown of all cohorts included in the mega-analysis, and their conclusions on sex bias on social motivation. Since these cohorts included groups of various genotypes and drug manipulations, only the untreated controls were included in the mega-analysis to test for baseline sex differences in social motivation. The FCGxMyt1L cohort described in this paper is included as 041723. For the FCGxMYT1L cohort, only mice with congruent gonadal/chromosomal sex were included in the analysis (XX3*^Sry-^* and XY*^Sry-^*3*^Sry+^*).

We first examined three major variables (mean rewards, mean interactions, and mean distance traveled), to determine if there were consistent effects of Sex, Age, or Cohort on social behavior across experiments. When examining social rewards, results from combining all social operant data revealed that cohort was the most significant single factor affecting variation in social seeking, with no effect due to sex (**Fig.S4)**. Cohort also was a highly significant factor in mean distance traveled, with no significant effect of sex **(Fig.S4)**. While there was no overall sex effect in distance traveled, cohort and sex seemed to significantly interact, with two cohorts demonstrating a strong female bias towards increased activity **(Fig.S4)**. Unlike rewards and interactions, only distance traveled showed a significant effect due to age, primarily driven by an increase in the 90-110 day group compared to the 44-62 day group **(Fig.S4)**. Collectively combined analysis of mean rewards and mean distance traveled indicated that from cohort to cohort, mice had substantial differences in both their drive for reward and total locomotion. This could be due to individual mouse variation or as consequence of adjustments to the social operant protocol over time.

For time in interaction zone (social orienting), there were significant effects of cohort **(Fig.S4)**, as seen in rewards and activity. Notably, this was the only variable to demonstrate a sex effect, with males spending more time in the interaction zone **(Fig.S4)**, Thus, males showing high social orienting appeared robust across most cohorts, although not seen in our model of chromosomal and gonadal sex effects. Since multiple factors were different between all cohorts, including litter composition, operant environment, and experimental question, it is not possible to pinpoint one factor causing the variation seen in social motivation across cohorts. Yet it is surprising how frequently a sex bias in mean interaction for individual cohorts appeared, many of which were properly powered for determining sex effects. Sex factors may be interacting with age, or some other unknown including baseline individual variation, to alter social motivation across cohorts.

## Discussion

In this study, we used the FCG mouse model to tease apart the impacts of chromosomal and hormonal sex on social behavior, both at baseline and in the context of genetic risk for a monogenic neurodevelopmental condition, MYT1L Syndrome. The data in this cohort revealed a strong bias in social motivation with the presence of ovaries, and by proxy female hormones, driving higher social seeking and orienting behaviors. Interestingly, while chromosomal sex did not appear significant in driving social motivation at baseline, in the context of MYT1L haploinsufficiency having two XX chromosomes led to increased social seeking and orienting. The sex by genotype interactions were not driven by differences in overall activity levels, as MYT1L mutation drove higher activity independent of chromosomal or gonadal sex. Combined, our behavioral data suggests complementary mechanisms for the sex bias in social behaviors, and a compensatory effect of X chromosome genes in the context of altered neurodevelopment.

Sex differences have long been observed in mammalian behavior, and classic interpretations of sex factors often assume the impact of sex chromosomes on output were mediated through gonadal hormones, since sex chromosome karyotype determines gonadal sex. However, while the *SRY* gene on the Y chromosome is the testes-determining factor leading to male gonad development, there are numerous other genes on the X and Y chromosomes that impact sex variation in biology independent of *SRY*[27]. Therefore, it is crucial when discussing sex differences to account for independent and potentially disparate effects of gonadal and chromosomal sex. This is especially true for neurodevelopmental conditions like autism, which disproportionately affect people with disorders of sexual differentiation[18,28], suggesting an interaction between sex factors (like hormones) and autism related traits. In addition, autism is 2-4 times more likely to be diagnosed in males[29] and a portion of this sex bias can be contributed to sex differences in heritability and genetic variance, but this sex ratio and core symptoms change with age and especially around puberty[30], highlighting the interplay between chromosomal and hormonal sex in complex behavior.

When comparing our results to prior work, animal behavior literature has significant evidence for female rodents being more active than males[9,31], including models of neurodevelopmental disorders (NDD)[32]. Higher female activity was seen in our previous MYT1L paper[16] and replicated in the FCGxMYT1L cohort during social operant and open field tasks. However, when separating chromosomal and gonadal sex using the FCG model, the sex factor driving higher female activity were different in both tasks. In the open field task, with no additional stimuli to capture the test subject’s attention, chromosomal sex seemed to drive female hyperactivity. In the context of a social stimuli in the operant task, gonadal sex drove the increased activity in females. The open field and social operant tasks are run for a similar length of time, but differ in environmental context, as mice in the open field are alone in a novel environment while mice in social operant are aware of a stimulus mouse in the other chamber. In addition, the distance traveled in the social operant task is an average across several days of 1-hr sessions, whereas the open field is one 1hr session. These differences in tasks could explain the varying effects of chromosomal vs. gonadal sex. It is possible that the novelty of the environment plays a role in the sex bias in activity seen in the social operant. We found gonadal sex to be a main driver of increased social orienting and seeking in the social operant task, and this increased social motivation likely contributed to increased activity, potentially hiding more subtle effects of chromosomal sex as seen in the open field task. Ovaries were associated with increased activity during both habituation and FR1 periods of the social operant task, suggesting this effect on locomotion is independent of task learning and instead attributable to the presence of a novel stimulus partner. Other studies evaluating the role of sex factors on activity using the FCG model have similarly found gonadal sex-driven higher activity in females[33], with some evidence for the role of the X chromosome in altering activity in the context of environmental manipulation[21]. Unlike rodents, human females typically tend to be less active than men, with weak evidence for changes in activity levels over hormonal cycles, such as menstruation[34]. It is of note that the largest study of activity across the menstrual cycle did find a correlation between increased physical activity and the luteal phase (high progesterone)[35], which coupled with anecdotal evidence suggests a role of sex hormones in activity. Yet larger comprehensive studies needed to truly determine the contribution of individual hormonal surges.

Here, both tasks support higher activity in female rodents generally, but in neither task did sex factors interact with MYT1L genotype. The influence of sex on increased activity seems more complex, as another *Myt1l* study found male-specific sex by genotype effects in activity. Overall, results from behavioral testing indicate that *Myt1l* mutations influence increased activity, as seen in previous studies[16,36,37]. This hyperactive phenotype complements the high rates of ADHD diagnosis in patients with MYT1L Syndrome[13] and suggests this effect may be independent of sex. As MYT1L Syndrome is a rare condition, it is not yet known if rates of ADHD co-morbidity have a similar male bias as seen in primary ADHD diagnosis[38]. Components of the open field, namely time spent into the center vs. perimeter, have been used to infer impacts on anxiety-like behavior. In our cohort, XY karyotype increased time spent in the center, suggesting a decrease in anxiety related behaviors compared to XX karyotype. However, open field on its own does not accurately encompass all aspects of anxiety-like behavior, and further evaluation through assays like the elevated plus maze are needed to follow up on this result.

Across tasks of social seeking and orienting in the social operant assay, gonadal sex was the primary driver of increased social motivation in females, regardless of MYT1L genotype. The impact of sex hormones such as estrogen and progesterone on sociability are complex, influenced by social task and brain region/method of manipulation, with estrogen potentially leading to increased social behaviors acutely[39,40]. The effect of estrogen in human behavior is less known, with some weak correlation between increased estrogen concentration and aggressive behavior[41]. On the other hand, testosterone in both rodents and humans has been found to increase behaviors associated with aggression and dominance, indirectly leading to decreased social behaviors[42]. Therefore, our results are consistent with what is known about the hormonal contribution to sex variation in social behavior.

Less is known of the independent contributions of X and Y chromosomes on social behavior in rodents. FCG studies have shown sex chromosome interactions in various behavior studies, including social behaviors[21]. These studies ultimately reveal the complex nature of sex chromosomal interactions, with effects varying by genotype, manipulation, and behavioral task. In the current study, chromosomal sex interacts with MYT1L genotype, driving higher social motivation in XX MYT1L Hets, relative to both XY MYT1L Hets and XX MYT1L WTs. These sex effects persist even when analyzing a subset of consistent achievers, implying the effect is not due to differences in task consistency or learning. Some effects seen in the full cohort were no longer significant in the consistent achiever subset, which we believe is due to decreased power per group. The pro-social effect of XX karyotype has been seen before in FCG studies. For example, use of FCG to investigate play behavior found XX sex chromosome karyotype drove increased social behaviors, and fewer exploratory/investigative behaviors[43]. As *Myt1l* is a neuronal transcription factor thought to act as either a repressor or enhancer depending on context[14–16], it is possible that MYT1L interacts with genes on sex chromosomes to mediate this sex by genotype effect. Single nuclei RNA sequencing data from MYT1L Het cortical samples reveals X chromosome genes that are differentially expressed by sex[44], and more investigation is needed to determine if MYT1L interacts with these gene sequences to facilitate sex by genotype effects.

Unexpectedly, the offspring proportions did not follow Mendelian inheritance patterns, with significantly fewer MYT1L heterozygote offspring in MYT1L x FCG litters. However, inheritance of the Y*^Sry-^* and 3*^Sry+^* chromosomes were as expected, in both FCG-only and FCGxMYT1L litters. *Myt1l* is a transcription factor on chromosome 2, making linkage and translocation with the *Sry* containing chromosome 3 impossible. Litters showed no differences in gross anatomy or ability to move, suggesting the interaction of MYT1L and FCG genotypes might cause lower *in utero* viability for MYT1L Het, regardless of sex factors. While there are no published studies reporting non-Mendelian inheritance of either FCG or MYT1L, it may be important for future researchers to consider viability when crossing the FCG genotype to other mouse models.

While our data does reveal distinct effects of gonadal and chromosomal sex, it is important to recognize the limitations of our interpretation. The mega-analysis of several social operant cohorts showed the sensitivity of the sex bias in social rewards, since directionality and effect size depended mostly on cohort. Machine learning analysis of spontaneous mouse behaviors have demonstrated individual variation as the main driver for differences in activity, even when accounting for hormonal state[45]. Therefore, this individual variation plus additional cohort effects could have hidden more subtle effects of sex. The social operant paradigm has evolved since first published, including alterations in light color and intensity, and apparatus flooring. In particular, the flooring changed from metal bars (associated with increased stress) to acrylic. Since stress responses contain significant sex variation, alterations of the environment such as this could contribute to variance in sex-specific social rewards across cohorts, as could differences in age[46]. In all, this retrospective mega analysis suggests some factors that deserve deliberate prospective studies to determine if they interact with sex.

Finally, recent information has revealed the FCG genotype mouse have additional unintended genetic modifications that may affect behavioral data interpretation. Specifically, a 3.2 MG region of the X chromosome was translocated to the Y*^Sry-^* chromosome, essentially making any of our XY*^Sry-^* groups have two copies of nine X chromosome genes[47]. Therefore, the FCG model in this case cannot give us a definitive case of X vs Y chromosome driving factors but is still valuable to separate chromosomal and gonadal effects. It is also possible these additional X chromosome genes in our Y caries may explain why males in this cohort did not show the robust increase in social orienting seen across the majority of the cohorts in the mega-analysis.

### Conclusions

This study is one of the first to use the FCG mouse model to tease apart mechanisms of sex by genotype effects in the context of neurodevelopmental disruption, demonstrating the independent contributions of sex chromosomes to behavioral changes in a MYT1L human variant model. MYT1L is not the only autism related gene to show differential sex effects, so similar experiments should be done in other models (*Shank3, Ube3a)* to determine how generalizable these gonadal and chromosomal contributions are to neurodevelopmental traits, and with updated FCG mice. Ultimately, understanding the mechanisms behind sex variation in behavior can help us better understand the basis of neuropsychiatric conditions with sex bias in prevalence and presentation. In addition, it ensures sex differences research is inclusive of varying expressions of sex, with the main goal of improving support options for all people with neurodevelopmental challenges.

## Acknowledgements

We would like to thank the members of the Dougherty Lab for guidance throughout the experimental process, Dr. Josh Rubin for providing the four-core genotype mice, and Kyle Kniepkamp for assistance designing and creating custom chambers. This work was generously supported by the NIH [T32GM007200-50 (MSTP@WUSM), RO1MH107515, R01MH124808] and Simons Foundation Autism Research Initiative (“Integrate Analysis of Mechanisms Underlying Sex Epistasis in Autism” to J.D.D).

---

**Supplementary Figure 1:**
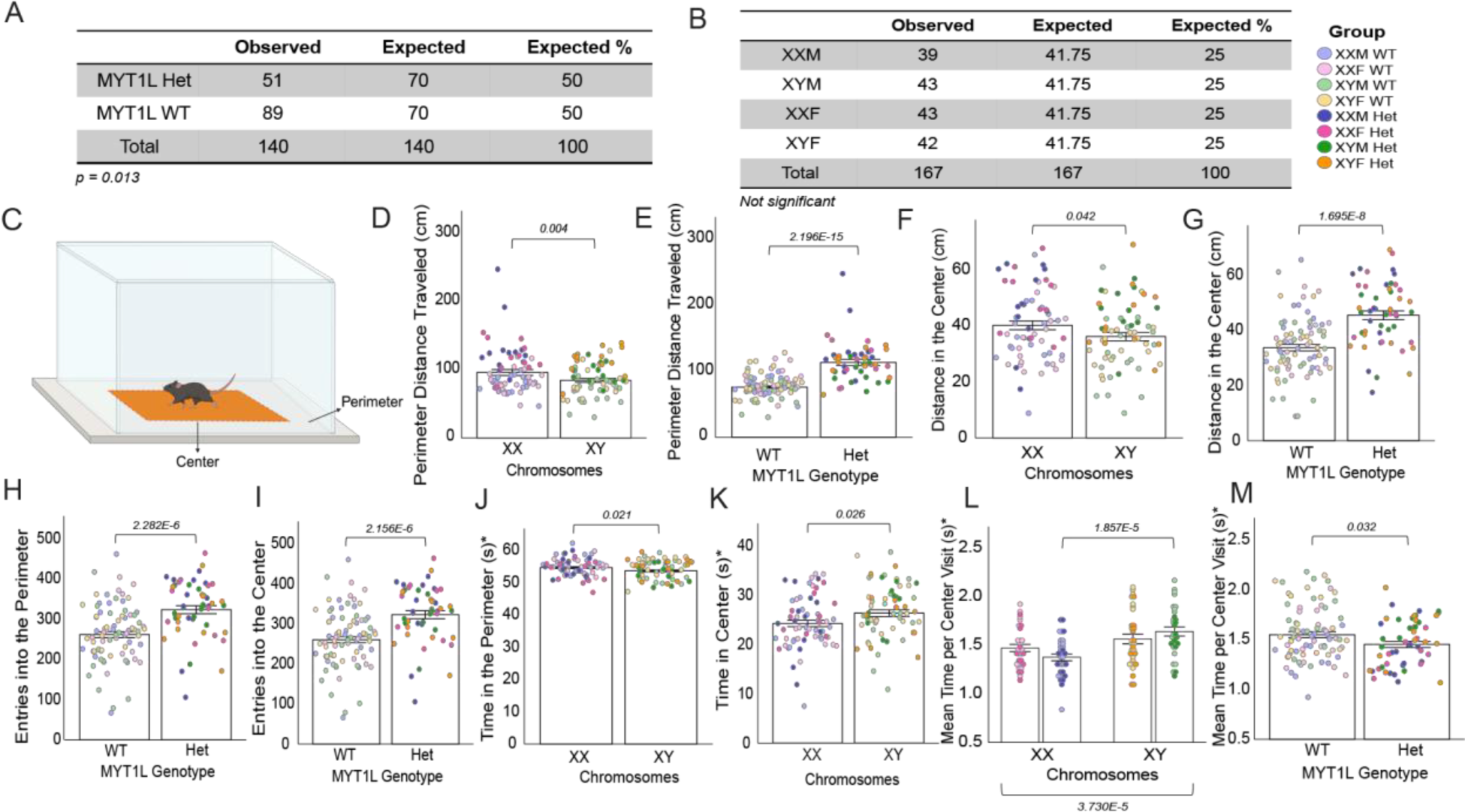
Open field assay revealed chromosomal and MYT1L genotype effects on activity. **A)** MYT1L genotype of all mice across 19 *Myt1l^+/-^ XX x Myt1l^+/+^ XY^Sry-^3^Sry+^*litters, including experimental mice used in open field and social operant assays. MYT1L mutation was inherited by significantly fewer offspring than expected by mendelian heritability pattern s. **B)** Sex factor breakdown of all mice across 24 litters, including experimental mice used in open field and social operant assays. The 24 litters included 19 *Myt1l^+/-^ XX x Myt1l^+/+^ XY^Sry-^3^Sry+^* litters, and 5 *Myt1l^+/+^ XX x Myt1l^+/+^ XY^Sry-^3^Sry^*. Inheritance of modified third chromosome with *Sry* gene followed expected mendelian heritability. **C)** Diagram of open field chamber. Dashed red line designates boundary between center zone (orange) and perimeter zone (no color). **D)** XX mice travel a greater distance in the perimeter than XY mice. **E)** MYT1L Het mice travel a greater distance in the perimeter than MYT1L WT mice. **F)** XX mice travel a greater distance in the center than XY mice. **G)** MYT1L Het mice travel a greater distance in the center than MYT1L WT mice. **H)** MYT1L Het mice entered the perimeter significantly more than MYT1L WT mice. **I)** MYT1L Het mice entered the center significantly more than MYT1L WT mice. **J)** XX mice spend more time in the perimeter zone than XY mice. **K)** XX mice spend less time in the center zone than XY mice. **L)** XY mice spend more time on average per visit in the center zone compared to XX mice. In the full univariate model, chromosomes and gonads interact to influence mean time per visit in the center (*p=0.048)*. Specifically, XX mice with testes typically spent less time per visit in the center while XY mice with testes spent significantly more time per visit in the center. **M)** MYT1L Het mice spend less time in the center per visit compared to MYT1L WT mice. For all panels, error bars indicate SEM. Asterisk (*) indicates variables that underwent square root transformation to normalize data distribution.

**Supplementary Figure 2:**
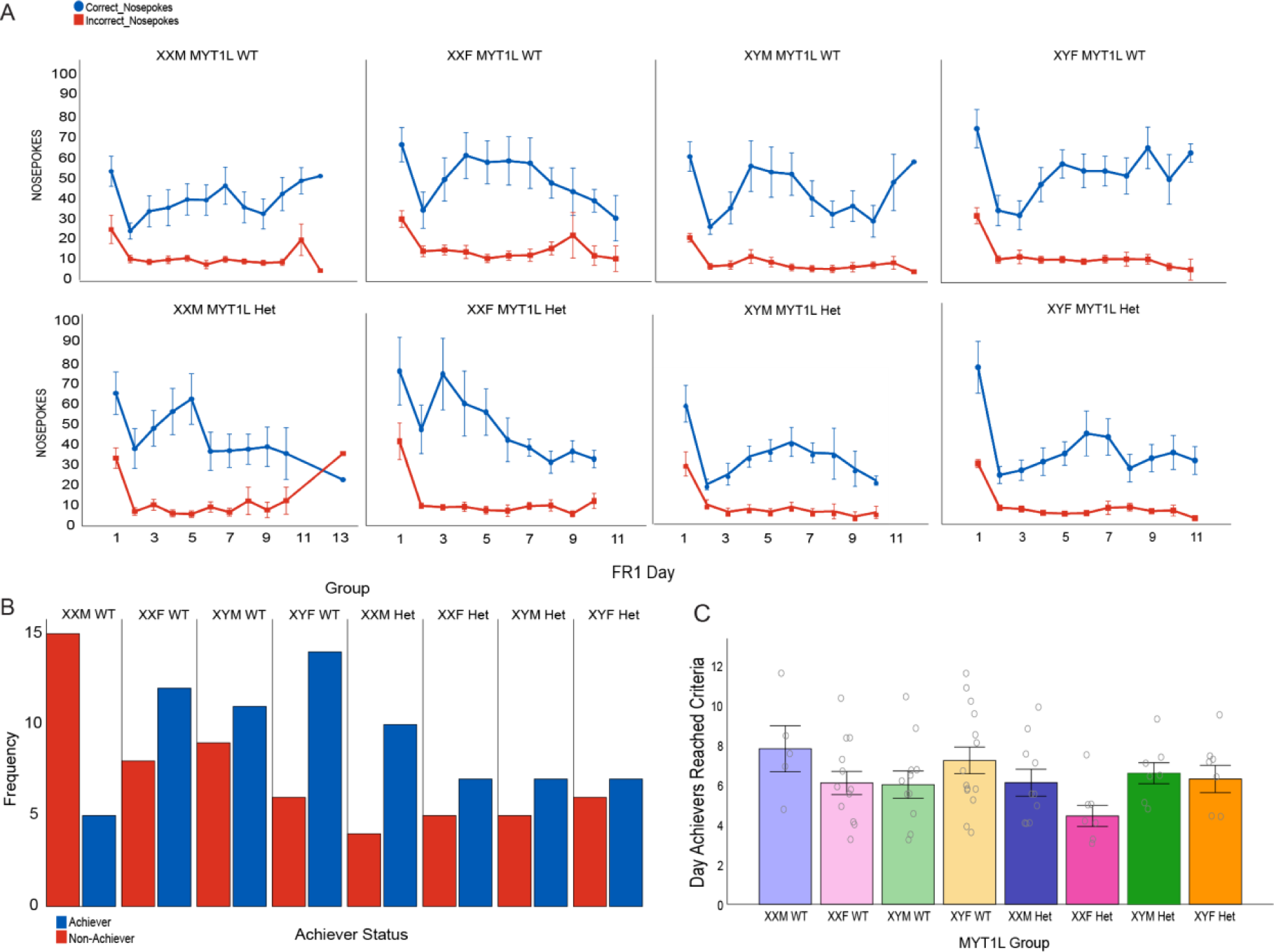
Conditioning achievement did not depend on MYT1L genotype or sex factors. **A)** Mean of daily nosepokes across the FR1 testing period, with correct nosepokes in blue and incorrect nosepokes in red. Top row are MYT1L WT and bottom row are MYT1L Het **B)** Histogram showing frequency of achievers (blue) and non-achievers (red) per group. **C)** FR1 day achievers reached the third consecutive day of criteria across the eight experimental groups. For all panels, error bars indicate SEM.

**Supplementary Figure 3:**
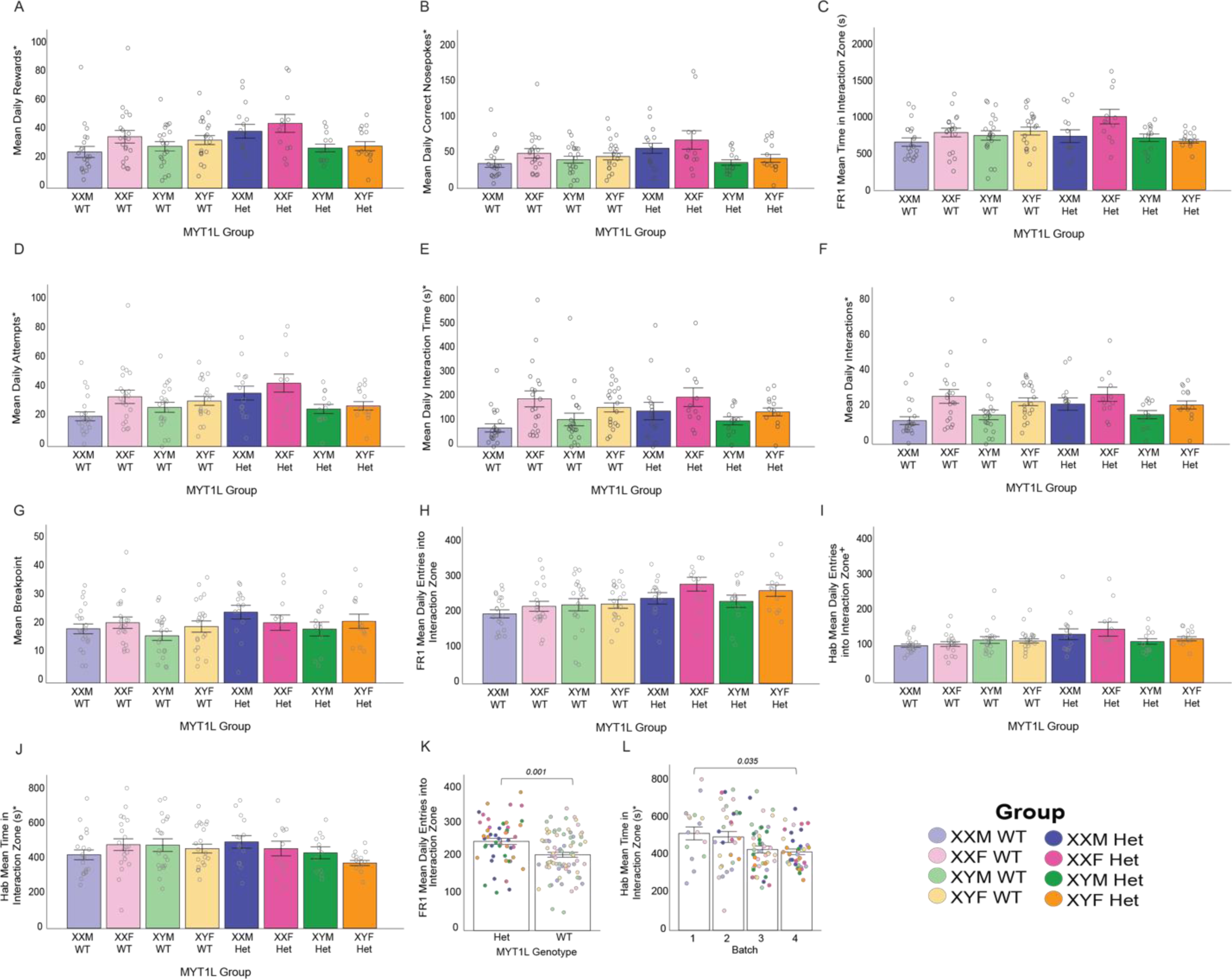
Gonadal and chromosomal sex independently act to alter social seeking and orienting in four core genotype mice. **A)** Mean Daily Rewards across all 8 groups. **B)** Mean Daily Correct Nosepokes across all 8 groups. **C)** Mean Time in the Interaction Zone across all 8 groups during FR1. **D)** Mean Daily Attempts across all 8 groups. **E)** Mean Daily Interaction Time across all groups. **F)** Mean Daily Interactions across all 8 groups. **G)** Mean breakpoint across all 8 groups, averaged from 3 separate PR trials. **H)** Mean Daily Entries into the Interaction Zone across all 8 groups during FR1. **I)** Mean Daily Entries into the Interaction Zone across all 8 groups during habituation. **J)** Mean Time in the Interaction Zone across all 8 groups during habituation. **K)** MYT1L Het test mice enter the interaction zone more often than MYT1L WT mice during FR1. **L)** Batch was the main driver of variation in Mean Time in the Interaction Zone during habituation, primarily driven by batch 4. For all panels, error bars indicate SEM. Asterisk (*) indicates variables that underwent square root transformation to normalize data distribution. Cross (^+^) indicates variables that underwent natural log transformation to normalize data distribution.

**Supplementary Figure 4:**
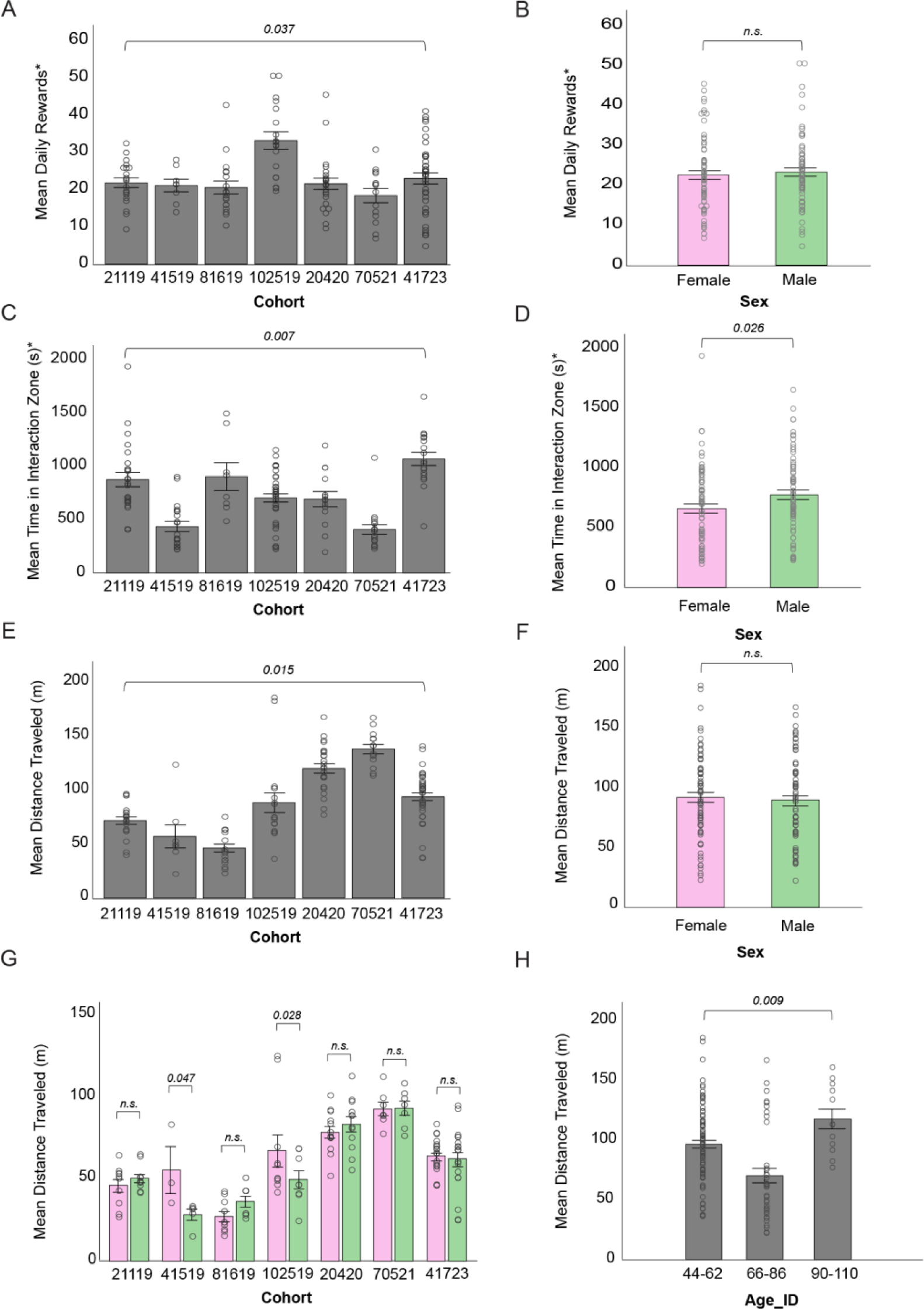
Mega-analysis shows variation in sex bias in social motivation is attributable to cohort effects. **A)** Mean Daily Rewards for control groups across 7 social operant cohorts, with cohort effect shown**. B)** Mean Daily Rewards for 7 social operant cohorts split by sex shows no difference in social seeking. **C)** Mean Time in the Interaction Zone with cohort effect shown**. D)** Mean Time in the Interaction Zone Rewards for 7 social operant cohorts split by sex shows males spend more time in the interaction zone than females. **E)** Mean Distance Traveled with cohort effect shown**. F)** Mean Distance Traveled in the Interaction Zone Rewards for 7 social operant cohorts split sex shows no difference in activity. **G)** Mean Distance Traveled for each social operant cohort split by sex demonstrates two social operant cohorts (041519 and 102519) show significantly higher activity in females than males. **H)** Mean Distance Traveled was the only variable to have significant effects with age, namely driven by the 66-86 age group. Statistics indicate results from cohort subset of mega-analysis ANOVA and multiple comparisons (**Table S5)** and is not the same as reported sex bias from original published experiments **(Table S4)** which includes all experimental groups. For all panels, error bars indicate SEM. Asterisk (*) indicates variables that underwent square root transformation to normalize data distribution before analysis.

**Supplementary Table 1:**

| Operant Interval | Social Motivation Component | Outcome | Definition |
| --- | --- | --- | --- |
| FR1 | Social Seeking | Conditioning Criteria Achieved | Animal showed conditioning to reward stimulus for three consecutive days (40 correct nosepokes, 75% poke accuracy, and 75% successful rewards) |
| FR1 |  | Reward | Door opening in response to correct nosepoke in FR1 |
| FR1 |  | Correct Nosepokes | Nosepoke into active nose hole, including during a reward |
| PR3 |  | Breakpoint | Maximum nosepokes performed to elicit a reward in PR testing |
| Hab, FR1 | Social Orienting | Total Time in Interaction Zone | Duration of time in which the test mouse is in the interaction zone, regardless of the stimulus mouse or whether the door is open |
| Hab, FR1 |  | Entries into Interaction Zone | Number of entries the test mouse makes into the interaction zone, regardless of the stimulus mouse or whether the door is open |
| FR1 |  | Attempts | Number of rewards where the test mouse is at the interaction zone with the door open, regardless of stimulus mouse |
| FR1 |  | Interactions | Number of rewards where test and stimulus mice are both at the interaction zone with the door open |
| FR1 |  | Interaction Time | Duration of time the test and stimulus mouse are simultaneously at the open door during a reward |

Adapted from Maloney et al. 2023[1]. Interaction zone depicted in Figure 2. Hab (Habituation); FR1 (Fixed Ratio 1); PR3 (Progressive Ratio 3)

**Supplementary Table 2:**
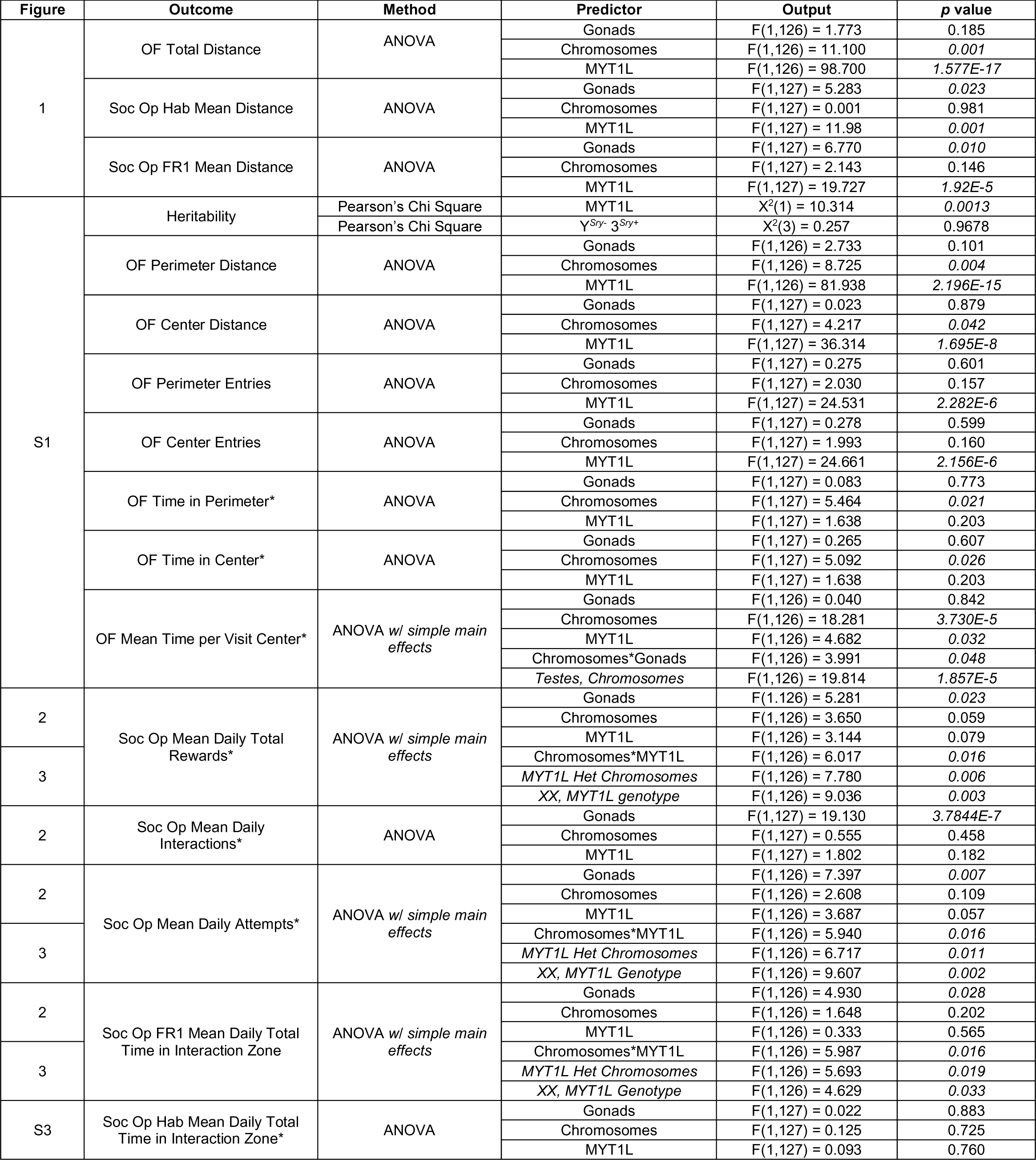
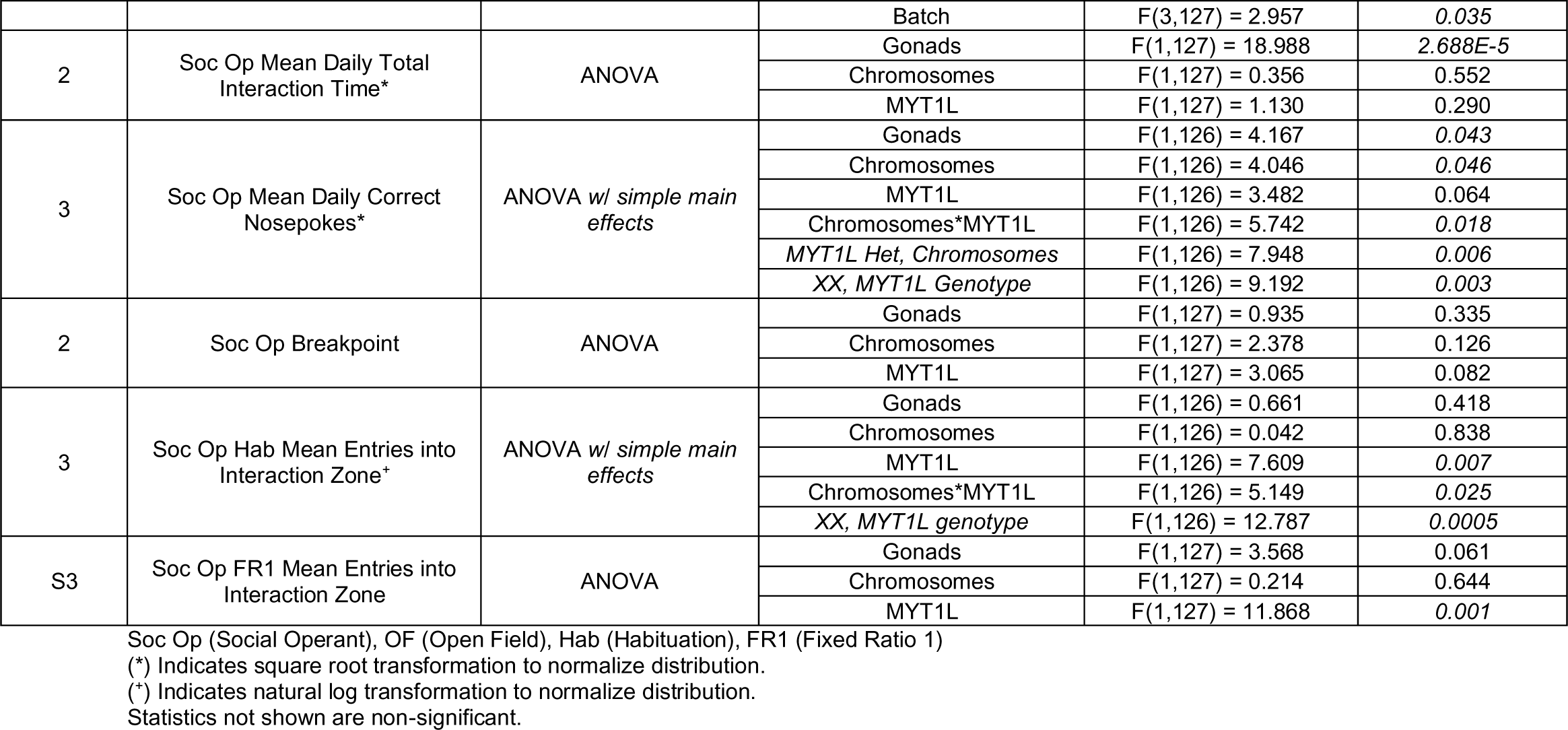

**Supplementary Table 3:**
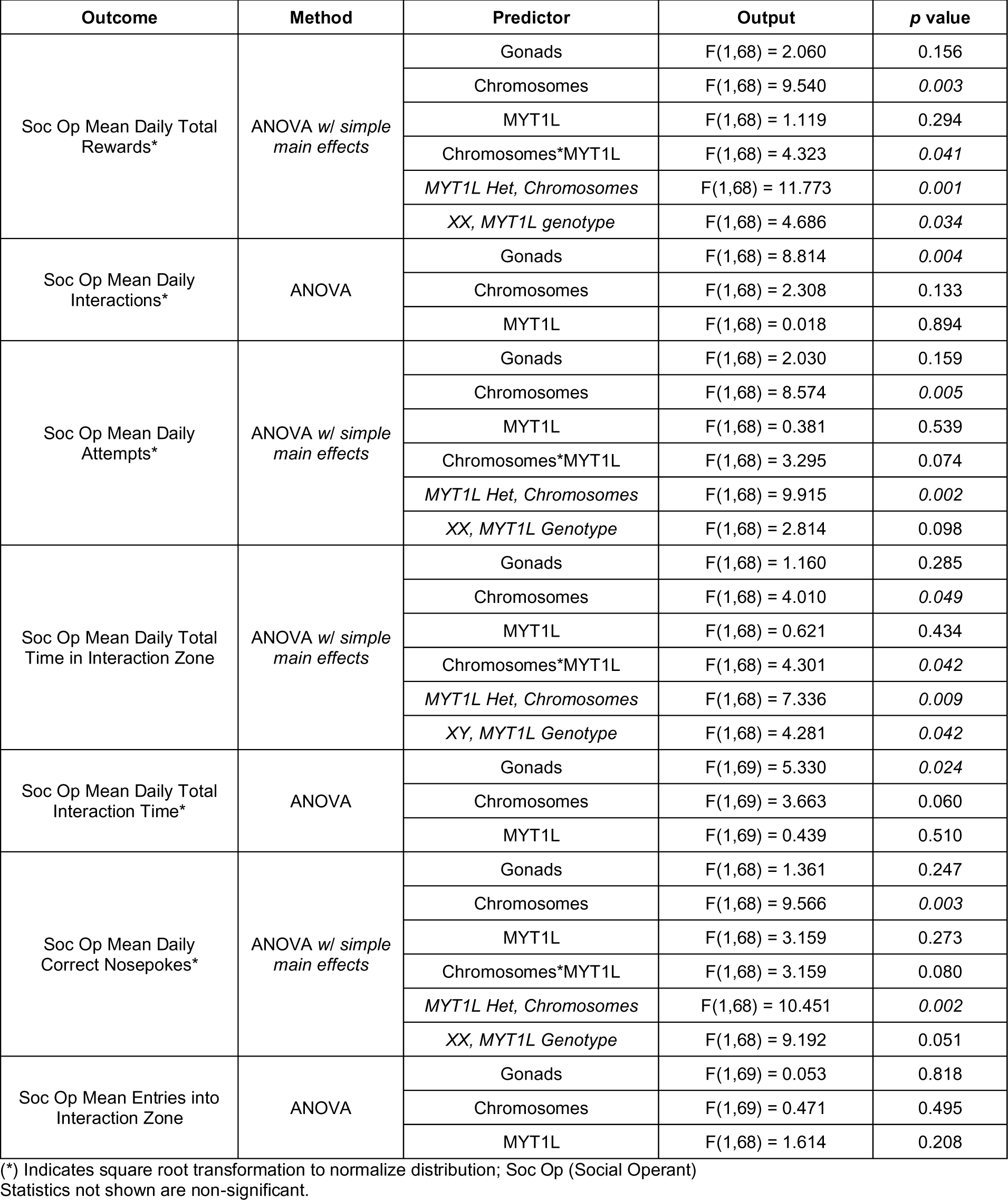
Analysis on subset of consistent achievers (Fig.S2B).

**Supplementary Table 4:** Summary of Cohorts Included in Social Operant Mega-Analysis.

| Cohort Number | Control Group | Age at start | Sample Size (M:F) | Sociability Bias in Original Analysis |
| --- | --- | --- | --- | --- |
| 021119 | WT C57 | P56-82 | 10:10 | M > F |
| 081619 | WT C57 | P70-99 | 8:10 | M = F |
| 041519 | WT C57 | P58-75 | 5:3 | M = F |
| 102519 | WT C57 | P47-50 | 8:10 | M > F |
| 020420 | WT CD1 | P40-97 | 12:13 | M = F |
| 070521 | WT C57 | P70-99 | 7:7 | M > F |
| 041723 | XY<sup>Sry-3<sup>Sry+</sup></sup>; XX3<sup>Sry-</sup> | P47-50 | 20:19 | M < F |

WT (Wildtype); M (Male); F (Female)

**Supplementary Table 5:** Mega-Analysis on 7 published social operant cohorts (Fig S3).

| Subset | Outcome | Method | Predictor | Output | p value |
| --- | --- | --- | --- | --- | --- |
| Combined Controls | Mean Daily Rewards* | ANOVA w/<br>cohort as<br>random factor | Sex | F(1,8.553) = 1.467 | 0.258 |
|  |  |  | Cohort | F(6,6) = 4.919 | 0.037 |
|  |  |  | Sex*Cohort | F(6,128) = 1.041 | 0.402 |
|  | Mean Total Time in Interaction Zone* | ANOVA w/<br>cohort as<br>random factor | Sex | F(1,7.409) = 7.693 | 0.026 |
|  |  |  | Cohort | F(6,6) = 9.719 | 0.007 |
|  |  |  | Sex*Cohort | F(6,129) = 1.861 | 0.092 |
|  | Mean Distance | ANOVA w/<br>cohort as<br>random factor +<br>simple main<br>effects | Sex | F(1,13.902) = 0.228 | 0.640 |
|  |  |  | Age | F(2,125) = 4.877 | 0.009 |
|  |  |  | 90-110 vs. 44-62 | 95% CI[905.129,4783.648] | 0.004 |
|  |  |  | Cohort | F(6,6) = 7.205 | 0.015 |
|  |  |  | Sex*Age | F(2,125) = 0.259 | 0.772 |
|  |  |  | Sex*Cohort | F(6,125) = 2.865 | 0.012 |
|  |  |  | 041519, Sex | F(1,125) = 4.021 | 0.047 |
|  |  |  | 102519, Sex | F(1,125) = 4.940 | 0.028 |
|  |  |  | Female, Cohort | F(6,125) = 16.877 | 3.336E-14 |
|  |  |  | Male, Cohort | F(6,125) = 10.788 | 1.195E-9 |

(\*) Indicates square root transformation to normalize distribution. Statistics not shown are non-significant.

